# Host-dependent salivary effector candidates and predicted plant immune targets in *Spodoptera frugiperda*

**DOI:** 10.64898/2026.09.21.753394

**Authors:** Sakshi Pandey, Sundaram Shilpi, Vineeta Roshanlal, Hanuman Sahay Meena, Chandra Pal Singh, Jayendra Nath Shukla

## Abstract

Salivary secretions of herbivorous insects contain proteins that modulate plant defence and facilitate herbivory. Despite the broad host range of *Spodoptera frugiperda*, a major agricultural pest, its salivary molecular arsenal involved in interactions with different host plants remains poorly understood. In particular, the salivary proteins induced during feeding on different host plants and their potential interactions with plant immune components remain largely unexplored.

To address this gap, we generated salivary gland transcriptomes from larvae reared on an artificial diet as well as larvae fed on tomato or chickpea plants. Comparative transcriptomic analysis identified salivary genes that were differentially expressed in response to plant feeding compared with the artificial diet. The encoded proteins were subsequently analysed using a secretory protein prediction pipeline to identify host-induced putatively secreted proteins with potential effector functions. Candidate salivary effectors were further investigated using large-scale *in silico* protein-protein interaction analyses with AlphaPulldown to predict their interactions with immune-pathway associated proteins of tomato and chickpea.

This analysis identified multiple putative salivary effector candidates whose expression was induced by plants feeding, with distinct candidate repertoires associated with tomato and chickpea. Predicted interactions between these candidates and plant immune-associated proteins further identified potential host targets. Together, these findings suggest that *S. frugiperda* may adjust its salivary molecular repertoire in response to the host plant and provide a transcriptome-wide framework for validating effector-target interactions and elucidating the molecular mechanisms underlying S. frugiperda-plant interactions.

## Introduction

Herbivorous insects are the major biotic constraints on crop production, and their interactions with host plants are shaped by reciprocal adaptations between insect feeding strategies and plant defence mechanisms. Herbivorous insects and their host plants have co- evolved over millions of years, resulting in complex interactions that shape their respective survival strategies. Plants have evolved constitutive and inducible defence mechanisms to combat insect attack, while insects have, in turn, developed strategies to overcome or manipulate these defences, thereby enabling successful feeding and colonization.

The oral secretions of herbivorous insects contain a diverse array of bioactive molecules, including herbivore-associated molecular patterns (HAMPs) and effector proteins, that play critical roles in plant-herbivore interactions. HAMPs, often referred to as elicitors, are recognised by plant receptors and can trigger jasmonate-mediated defence responses (Malik, Kumar, & Nadarajah, 2020). In contrast, effector proteins can target key plant signalling- pathways to modulate or suppress host defence, thereby facilitating insect feeding and development (Snoeck, Guayazán-Palacios, & Steinbrenner, 2022). Thus, insect saliva constitutes an important molecular interface between herbivores and plant immunity and represents a potential source of proteins involved in host adaptation.

Insects with piercing and sucking mouthparts inject saliva directly into plant tissues, whereas insects with chewing mouthparts release saliva at feeding site (H. Wang, Shi, & Hua, 2023). Although majority of studies of insect effectors have primarily focused on sap-feeding insects, increasing evidence demonstrates that salivary effectors from chewing insects also play important roles in suppressing plant defence. Research on effector from chewing insects began with the discovery of glucose oxidase (GOX) from the labial glands of *Helicoverpa zea* (Eichenseer, Mathews, Bi, Murphy, & Felton, 1999). GOX catalyses the oxidation of glucose to gluconic acid and hydrogen peroxide, which inhibits nicotine production, and modulates reactive oxygen species (ROS) response in tobacco. Subsequent studies have revealed salivary proteins from chewing insects and demonstrated their effects on plant defence signalling. HARP1 (*Helicoverpa armigera* R-like Protein 1), identified from *H. armigera*, stabilizes JAZ (Jasmonate ZIM-Domain) repressors and prevents their COI1-mediated degradation, thereby suppressing jasmonate signalling (C. Y. Chen et al., 2019). Similarly, Hexenal isomerase (Hi-1), a member of the GMC oxidoreductase family identified from the oral secretions of *Manduca sexta*, converts (Z)-3-hexenal into its (E)-2-hexenal isomer, thereby modulating plant volatile emissions and defence signalling (Lin et al., 2023). More recently, HSC70-3 from *Plutella xylostella* was shown to suppress host defence gene expression (Qiao et al., 2025), while HYPB1, from *Helicoverpa armigera* targets the cotton dirigent protein GhDIR15 and affects the biosynthesis of plant secondary metabolites (Y. Wang et al., 2026). Together, these studies indicate that chewing insects possess diverse salivary proteins capable of influencing multiple components of plant defence.

The fall armyworm, *Spodoptera frugiperda* (J.E. Smith), is a highly polyphagous lepidopteran pest with chewing mouthparts. It feeds on approximately 350 plant species belonging to 76 families, including maize, sorghum, sugarcane, rice, wheat, cotton, tomato and several other economically important crops, causing substantial yield losses worldwide (Kenis et al., 2023). Its broad host range makes *S. frugiperda* a useful system for investigating the molecular mechanisms that enables this invasive pest to colonize different host plants.

The remarkable ability of *S. frugiperda* to exploit diverse host plants is considered a key factor underlying its widespread ecological expansion. However, the molecular mechanisms underlying this host adaptability remains poorly understood, particularly the role of salivary proteins in modulating host plant defence. Recent studies have begun to reveal the effector potential of *S. frugiperda* saliva. A proteomic study identified a Venom R-like protein (VRLP4) in *S. frugiperda* oral secretions that suppresses defence-related gene expression in wounded leaves (Xian Zhang et al., 2024). Another salivary protein, protein disulfide isomerase (SfPDI), was reported to function as a redox-active effector that modulates the folding and stability of host defence proteins, thereby influencing ROS accumulation and plant resistance responses (Cai et al., 2025). However, the extent to which the salivary repertoire of *S. frugiperda* changes in response to different host plants remains unclear. In particular, the host-dependent induction of salivary proteins and their potential interactions with components of plant immune pathways have not been systematically investigated.

To investigate whether host plant feeding is associated with changes in the salivary molecular repertoire of *S. frugiperda*, we generated salivary gland transcriptomes (sialotranscriptomes) from sixth-instar larvae reared on an artificial diet or fed on two economically important crop plants, tomato (*Solanum lycopersicum*) and chickpea (*Cicer arietinum*). Salivary genes upregulated in response to plant feeding were analysed using a secretory protein prediction pipeline to identify host-induced putatively secreted proteins with potential effector functions. Candidate effectors were subsequently examined through *in silico* protein-protein interaction analysis with immune-associated proteins of tomato and chickpea using AlphaPulldown (Homma, Lyu, & van der Hoorn, 2024). By integrating host-dependent transcriptomic responses, secretory protein prediction and predicted protein-protein interactions, we identified multiple salivary effector candidates associated with different host plants and their potential plant immune targets.

## Materials and methods

### Plant growth conditions, insect rearing, and experimental feeding setup

Seeds of tomato (*Solanum lycopersicum,* cv. Arka vikas) and chickpea (*Cicer arietinum*, RSG-888) plants were procured from the Indian Institute of Horticultural Research, ICAR, Bengaluru, India, and the Rajasthan Agricultural Research Institute, ICAR, Jaipur, India, respectively. Seeds were surface sterilised using 4% (w/v) Sodium hypochlorite solution and directly sown in plastic pots (6.5 cm diameter x 8.5 cm height) containing 1:1 mixture of soilrite and perlite. Plants were maintained in a growth chamber (SR Lab instruments, India) at 22°C during the light period and 16°C during the dark period, with 70% relative humidity and a 15 h light:9 h dark photoperiod until they reached the vegetative stage, corresponding to approximately 30 days after sowing for tomato and 25 days after sowing for chickpea.

Eggs of *Spodoptera frugiperda* were procured from the National Bureau of Agricultural Insect Resources (NBAIR), Bengaluru, India. Larvae were reared under controlled conditions at 26 ± 1 °C, 60-70% relative humidity, and 15 h light:9 h dark photoperiod. Larvae were maintained on an artificial diet described by Shilpi et al. (2025) until sixth instar stage.

Same age of sixth instar larvae were randomly assigned to three feeding treatments: artificial diet, tomato and chickpea. Each treatment comprised of 30 larvae. Sixth instar larvae in the plant-feeding treatments were transferred individually to 30-day-old tomato or 25-day-old chickpea plants, with one larva maintained per plant. Larvae assigned to the control treatment were maintained on the artificial diet. Following 12 h of feeding, larvae from all three treatments were collected for salivary gland dissection. The experimental design therefore comprised three feeding conditions: artificial diet-fed (AD), tomato-fed (TP) and chickpea-fed (CP) larvae.

### Salivary gland collection and RNA extraction

Larvae were dissected individually in sterile 1x phosphate-buffered saline (PBS) under a stereo microscope (Leica S9i). Salivary glands were carefully removed, immediately flash-frozen in liquid nitrogen and stored at −80 °C until RNA extraction. Two independent biological replicates were prepared for each feeding condition, with each replicate consisting of salivary glands pooled from 15 larvae.

Total RNA was extracted from frozen salivary gland samples using TRIzol reagent according to the manufacturer’s protocol. RNA quality was assessed by agarose gel electrophoresis, and RNA concentration was measured using a NanoQ spectrophotometer (KLAB Technology).

### Library preparation and RNA sequencing

RNA samples were sequenced by Eurofins Genomics (Bengaluru, India). RNA integrity was assessed prior to library preparation. Strand-specific paired-end RNA-sequencing libraries were prepared using the NEBNext Ultra II Directional RNA Library Prep Kit for Illumina (New England Biolabs, USA), following the manufacturer’s instructions.

Briefly, mRNA was enriched from total RNA using oligonucleotide-conjugated magnetic beads and subsequently fragmented enzymatically. First strand cDNA was synthesized using the NEBNext First Strand Synthesis Enzyme Mix (NEB), followed by second-strand cDNA synthesis using the NEBNext Second Strand Synthesis Enzyme Mix. Double stranded cDNA was purified using AMPure XP beads and subjected to A-tailing and adapter ligation. Adapter-ligated libraries were amplified by a limited number of PCR cycles and subsequently purified using AMPure XP beads. Library quality and fragment-size distribution were assessed using a 4200 TapeStation system (Agilent Technologies) with high-sensitivity D1000 ScreenTape. Libraries were sequenced on an Illumina sequencing platform using 2 × 150 bp paired-end chemistry.

### Processing and alignment of RNA-sequencing reads

Raw paired-end sequencing reads were quality-filtered using Trimmomatic v0.39. Adapter sequences, reads containing more than 5% ambiguous nucleotides and reads in which more than 10% of bases had a Phred quality score below 25 were removed. The resulting high-quality paired-end reads were used for downstream analyses.

The *S. frugiperda* reference genome and corresponding gene annotation files were obtained from the Ensembl Metazoa database (release 60). Quality-filtered reads from each biological replicate were aligned independently to the reference genome using STAR v2.7.10a with default parameters. The resulting alignment files were used for read quantification and differential gene-expression analysis.

### Gene-expression quantification and differential expression analysis

Gene-level read counts were generated from the STAR alignment files using featureCounts v2.0.3, with uniquely mapped reads assigned to annotated genes. Differential expression analysis was performed in R using the DESeq2 package. To identify salivary transcriptional responses associated with host-plant feeding, transcriptomes from tomato-fed and chickpea- fed larvae were each compared with the artificial diet-fed control group. Genes showing positive log2 fold-change values were classified as upregulated and genes showing negative log2 fold-change values as downregulated. Statistical significance was assessed using a p-value threshold of 0.05.

For visualization of expression patterns, normalized expression values were log-transformed and subjected to hierarchical clustering using Pearson uncentered distance and average-linkage clustering. Heat maps were generated from the resulting expression matrix.

### Transcriptome validation

To validate the transcriptome data, two genes from the tomato- and chickpea-fed transcriptomes were selected using stringent criteria: log2 fold change >1 and a base mean expression value >50 to ensure both biological significance and adequate expression levels. Primers for the selected genes were designed using Primer-3 software (https://bioinfo.ut.ee/primer3-0.4.0/).

Infestation assays were conducted again following the same protocol as the feeding experiment. Total RNA was extracted from pooled salivary glands using the TRIzol reagent (Invitrogen, India) following the manufacturer’s instructions. RNA samples were subsequently treated with DNase (Promega) to remove genomic DNA contamination. The concentration and purity of DNA-free RNA samples were determined by using the NanoQ Series (KLAB Technology). 3 μg of RNA per sample was reverse transcribed using a 17- mer oligo-dT primer and GoScript reverse transcriptase (Promega) enzyme for the synthesis of first-strand cDNA. Quantitative real-time PCR (qRT-PCR) was performed using Luna Universal qPCR Master Mix (New England Biolabs) in a CFX96 touch real-time PCR detection system (Bio-Rad) with the cycling parameter of initial denaturation at 95°C for 30 s, followed by 40 cycles of 95°C for 15 s, 60°C for 45 s and 72°C for 5s.

Relative expression levels were normalized against the endogenous control gene ribosomal protein 10 (RPL-10) of *S. frugiperda*. Data analysis was carried out using the 2^–ΔΔCT method, and statistical significance between tomato-fed, chickpea-fed, and control larvae was evaluated using an unpaired Student’s t-test.

### Functional enrichment and pathway analysis

Gene Ontology (GO) enrichment and Kyoto Encyclopedia of Genes and Genomes (KEGG) pathway analyses were performed using the ClusterProfiler R package (v4.2.2). GO enrichment was used to identify biological processes, molecular functions and cellular components overrepresented among differentially expressed genes. KEGG pathway annotation was performed by mapping *S. frugiperda* genes to insect pathways using the KEGG Automatic Annotation Server (KASS).

### Prediction of putatively secreted salivary proteins

Protein-coding sequences corresponding to transcripts upregulated in tomato-fed and chickpea-fed larvae were retrieved from the NCBI Protein database. The encoded proteins were screened using an *in silico* secretory-protein prediction pipeline based on SignalP, TMHMM, TargetP and WoLF PSORT, following the general strategy described by Shilpi et al. (2025).

SignalP was used to identify proteins containing an N-terminal signal peptide indicative of entry into the secretory pathway. TMHMM was used to predict transmembrane helices, and proteins containing predicted transmembrane regions were excluded from further analysis. TargetP was used to identify proteins predicted to contain mitochondrial targeting peptides; proteins with mitochondrial targeting predictions were also excluded from further analysis. WoLF PSORT was subsequently used to assess predicted extracellular localization.

Proteins satisfying the combined criteria of a predicted signal peptide, absence of transmembrane regions, absence of a predicted mitochondrial targeting peptide and predicted extracellular localization were classified as putatively secreted proteins. Among these, proteins derived from transcripts upregulated following plant feeding were retained as putative salivary effector candidates for downstream protein-protein interaction analysis.

### Identification of plant biotic stress- and defence-associated proteins

Reference proteomes of tomato and chickpea were retrieved from UniProtKB using the *S. lycopersicum* proteome identifier UP000004994 and the *C. arietinum* proteome identifier UP000087171, respectively.

Plant proteins associated with biotic stress and defence responses were identified based on Gene Ontology annotations. The search included GO terms associated with immune response, salicylic acid signalling, jasmonic acid signalling, reactive oxygen species responses, defence responses to insects and other plant defence-related biological processes. The resulting protein sets were used as candidate host targets for protein-protein interaction prediction with the *S. frugiperda* putative effector candidates.

### Prediction of salivary effector-plant protein interactions

Potential interactions between putative salivary effector candidates of *S. frugiperda* and plant defence-associated proteins were predicted using AlphaPulldown v2.1.1 (Yu, Chojnowski, Rosenthal, & Kosinski, 2023). The analysis included 43 salivary secretory proteins identified from tomato-fed larvae and 51 salivary secretory proteins identified from chickpea-fed larvae, together with 55 tomato and 42 chickpea proteins associated with plant immune and defence responses.

AlphaPulldown was implemented through a command-line workflow for multimeric protein- structure and interaction prediction. Multiple sequence alignments (MSAs) for the input proteins were generated using HH-suite v3.3.0 and HMMER v3.4, including Jackhmmer and HHblits searches. Sequence and structural searches incorporated UniRef90, UniRef30, UniProt, BFD, PDB70, MGnify and SmallBFD databases. The resulting sequence alignments and template information were processed using the create_individual_features.py workflow to generate the feature and template files required for downstream modelling. Protein complexes were subsequently modelled using the multimer workflow implemented through run_multimer_jobs.py. All computations were performed in a Python 3.11 environment using JAX v0.4.1 and JAXLIB v0.4.1+cuda11.cudnn86.

### Refinement and structural analysis of predicted protein complexes

To evaluate the effect of signal peptides on predicted protein-protein interactions, the ten highest-ranked *S. frugiperda* secretory protein candidates from each plant-feeding condition were subjected to a second round of AlphaPulldown analysis after removal of the predicted N-terminal signal peptide sequences. The resulting complexes were evaluated using AlphaFold-derived confidence metrics, including interface predicted template modeling (ipTM), predicted aligned error (PAE) and predicted local distance difference test (pLDDT) scores.

The highest-confidence interaction pairs from the tomato- and chickpea-associated candidate sets were selected for structural interface analysis. Protein-protein interfaces were characterized using PDBePISA, from which interface area, interacting residues, hydrogen bonds, salt bridges and solvation free-energy gain (ΔG) were obtained. Predicted complexes and interaction interfaces were visualized using UCSF Chimera v1.19. A summary of the predicted complexes was compiled based on the corresponding confidence metrics, including ipTM, combined ipTM+pTM.

## Result

### Illumina sequencing data

A total of six RNA-seq libraries were generated from pooled salivary gland samples of *S. frugiperda* larvae reared under three dietary conditions: artificial diet (AD), artificial diet followed by chickpea plant feeding (CP), and artificial diet followed by tomato plant feeding (TP), with two biological replicates per condition. All six RNA samples isolated from pooled salivary glands exhibited good RNA quality and acceptable RNA integrity (RIN) scores. Sequencing yielded between 8.5 and 13.7 million raw reads per sample.

Following quality filtering to remove low-quality and adapter-contaminated reads, 7,183,569-12,617,819, high quality (HQ) reads were obtained per library, corresponding to 1.98-3.59 Gb sequence data per sample. Between 80.0% and 82.1% of the HQ reads were uniquely mapped to the *S. frugiperda* reference genome. The sequencing and mapping statistics are summarized in Table 1.

**Table 1:** Summary of RNA quality, sequencing output, and mapping statistics for salivary gland transcriptome libraries of *S. frugiperda* larvae reared on different diets.

| Diet type | Artificial Diet (AD) | Chickpea plant (CP) | Tomato Plant (TP) |
| --- | --- | --- | --- |

| Samples | *AD-1* | *AD-2* | *CP-1* | *CP-2* | *TP-1* | *TP-2* |
| --- | --- | --- | --- | --- | --- | --- |
| RIN value | 9.8 | 9.4 | 9.8 | 9.6 | 9.6 | 8.9 |
| Raw Reads | 1,34,49,351 | 1,14,33,465 | 1,21,78,398 | 1,37,52,256 | 84,96,804 | 1,34,47,920 |
| Clean Reads (HQ) | 1,21,51,943 | 1,02,90,018 | 1,11,57,947 | 1,26,17,819 | 71,83,569 | 1,15,78,981 |
| HQ Base Pairs (GB) | 3.43 | 2.92 | 3.16 | 3.59 | 1.98 | 3.23 |
| Average read length (bp) | 412 | 409 | 392 | 411 | 388 | 409 |
| Mapped Reads (%) | 81.73 | 81.17 | 80.66 | 80 | 81.77 | 82.09 |

### Differentially expressed genes in response to host-plant feeding and their validation using qRT-PCR

A comparative analysis of the salivary transcriptomes of *S. frugiperda* was conducted to evaluate transcriptional changes associated with feeding on tomato and chickpea plants. The salivary transcriptomes of TP- and CP-fed larvae were independently compared with those of larvae maintained on AD. The analysis identified 353 upregulated genes in TP fed larvae and 1,017 upregulated genes in CP-fed larvae (log2 FC > 0.5; Fig. 1; Supplementary Table S1). In contrast 420 and 860 genes were downregulated in TP- and CP-fed larvae, respectively (log2 FC < -0.5; Fig. 1; Supplementary Table S2).

**Figure 1.**
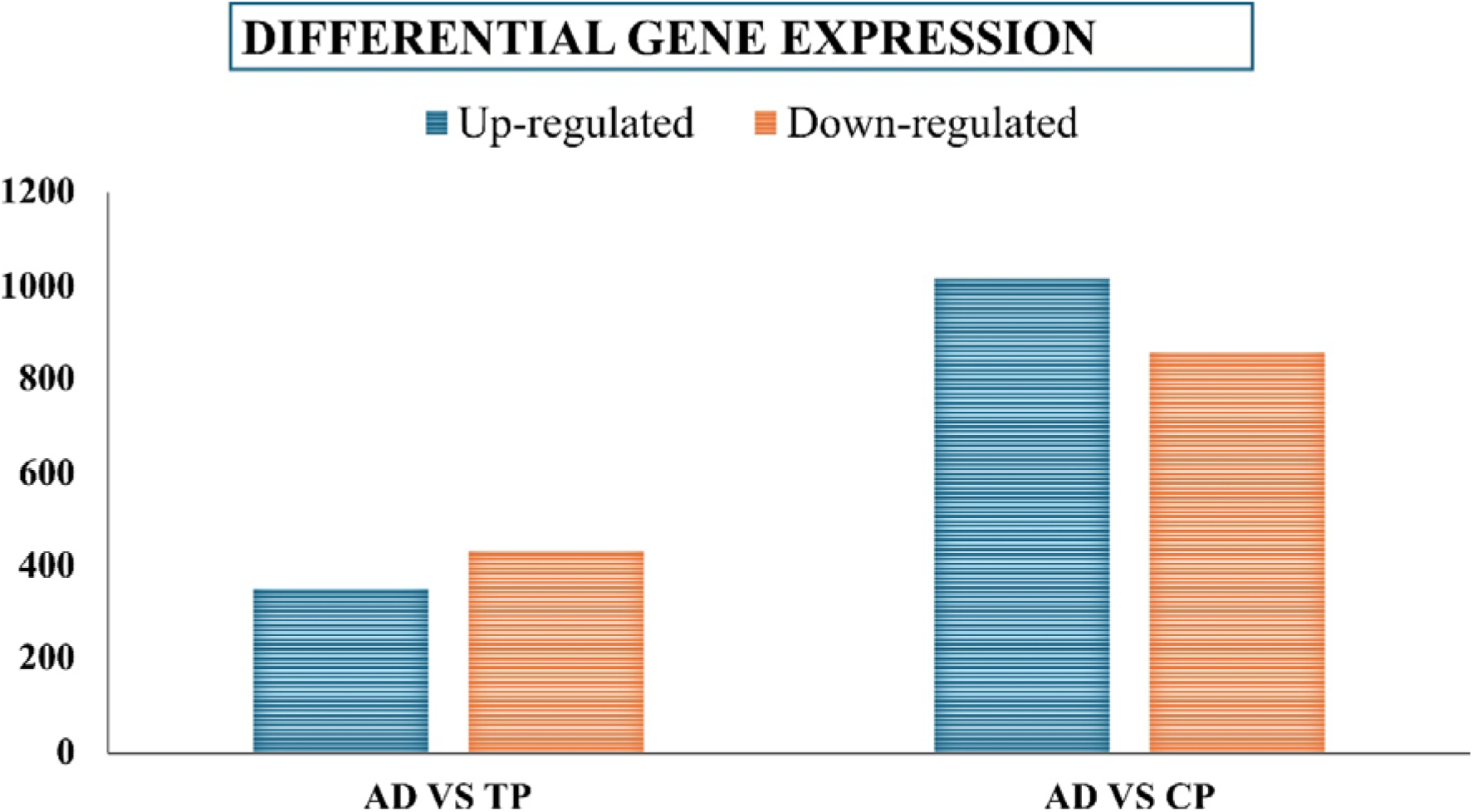
Bar graph representing the number of differentially expressed genes (DEGs) in salivary glands of larvae fed on tomato plants (TP) and chickpea plants (CP) relative to the artificial diet-fed control (AD). The Y-axis represents the number of differentially expressed genes.

The qRT-PCR results showed that the selected upregulated genes in tomato and chickpea-fed larvae of *S. frugiperda* showed upregulation pattern consistent to the trend observed in the transcriptomic analysis (Fig.2). The observed upregulation was statistically significant (*p* < 0.05). These results provided independent confirmation of the transcriptome data.

**Fig. 2.**
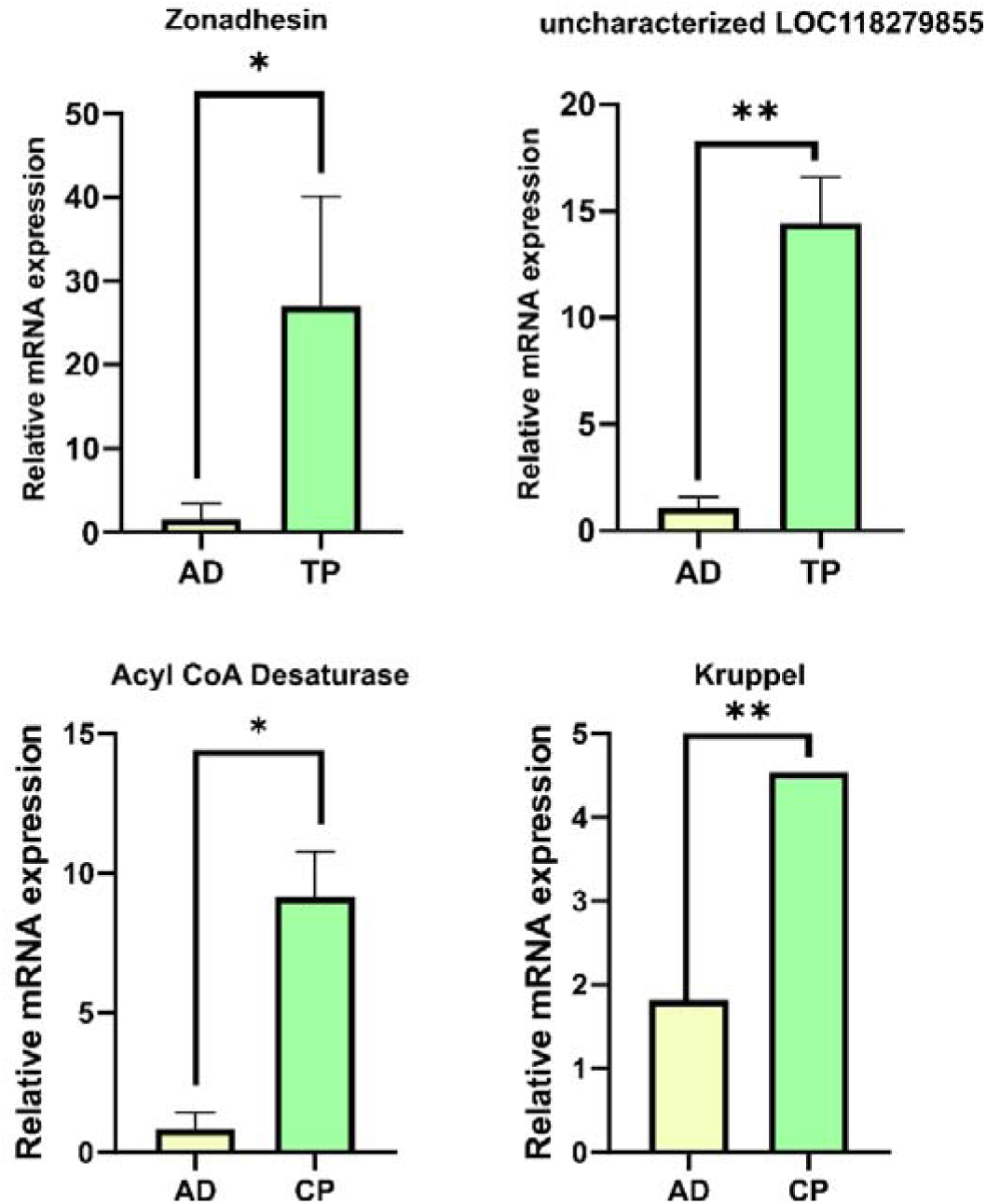
Relative mRNA expression levels of selected differentially expressed genes identified from the *S. frugiperda* salivary gland transcriptome were validated by quantitative real-time PCR (qRT-PCR). Zonadhesin and uncharacterized protein LOC118279855 were evaluated in salivary glands of insects fed on artificial diet (AD) and tomato plant (TP), whereas Acyl- CoA desaturase and Kruppel were evaluated in insects fed on artificial diet (AD) and chickpea plant (CP). Relative expression was calculated using the method after normalization to the reference gene (RPL-10). Bars represent mean relative expression ± SEM. Statistical analyses were performed using ΔCt values, with significance indicated by *P* < 0.05 (*).

To visualize differences in transcript abundance among the dietary conditions, heat-maps, scatter plots, and volcano plots were generated (Figs. 3 and 4). Hierarchical clustering of the top 50 DEGs showed distinct expression profiles between plant-fed and AD-fed larvae (Fig. 3C and 4C).

**Figure 3.**
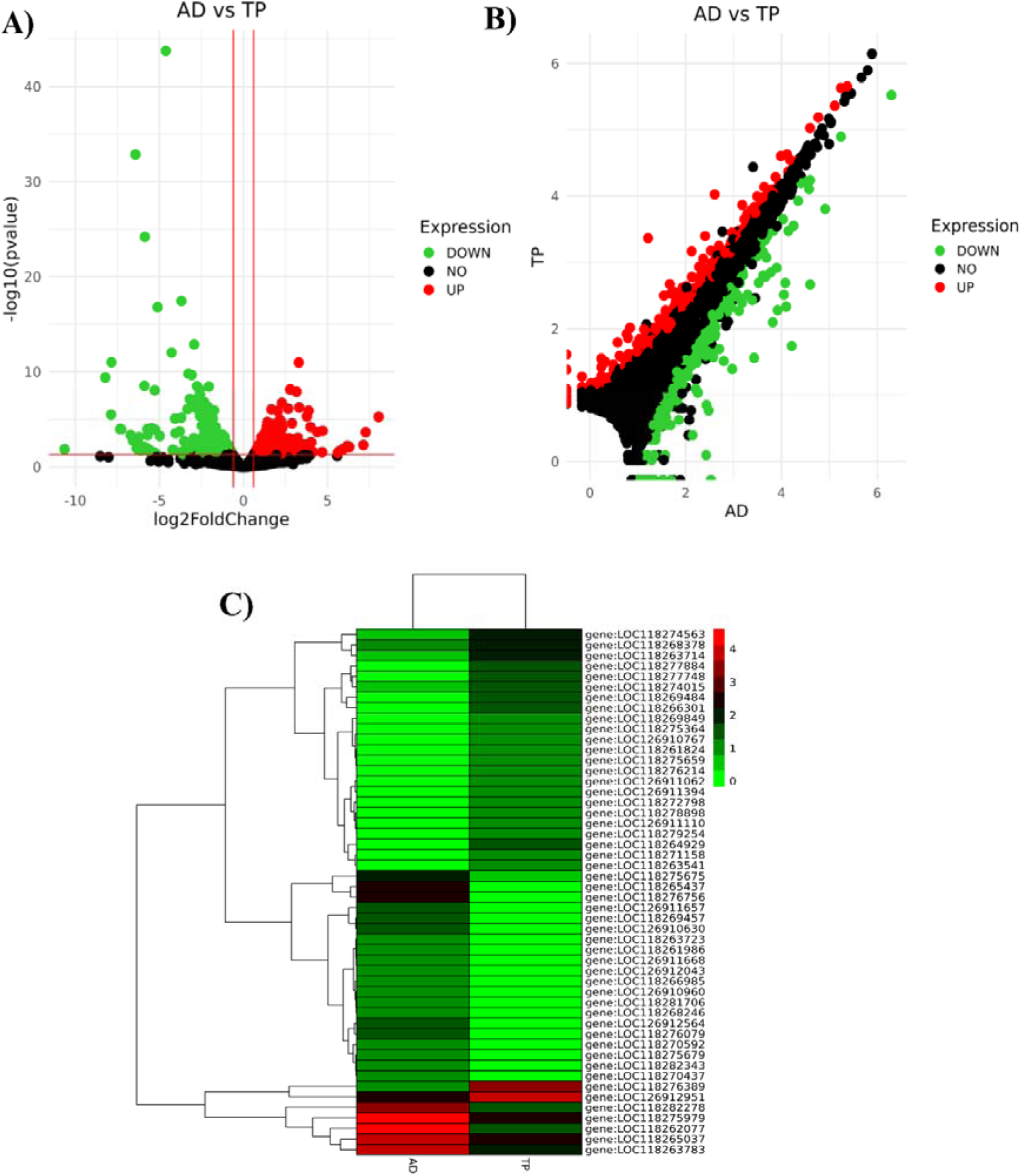
Transcriptome analysis of differentially expressed genes (DEGs) in the salivary glands of *Spodoptera frugiperda* larvae fed on tomato plants (TP) compared with those fed on artificial diet (AD). **A)** Volcano plot showing the distribution of significantly upregulated and downregulated genes based on log_2_ fold change and statistical significance (−log_₁₀_ *P*-value). Red dots indicate significantly upregulated genes, green dots indicate significantly downregulated genes, and black dots represent genes with no significant differential expression. **B)** Scatter plot showing the overall distribution of gene expression in TP- and AD-fed larvae. Red and green dots represent significantly upregulated and downregulated genes, respectively. **C)** Heat-map depicting the hierarchical clustering and expression patterns of selected DEGs, where red indicates higher expression and green indicates lower expression.

**Figure 4.**
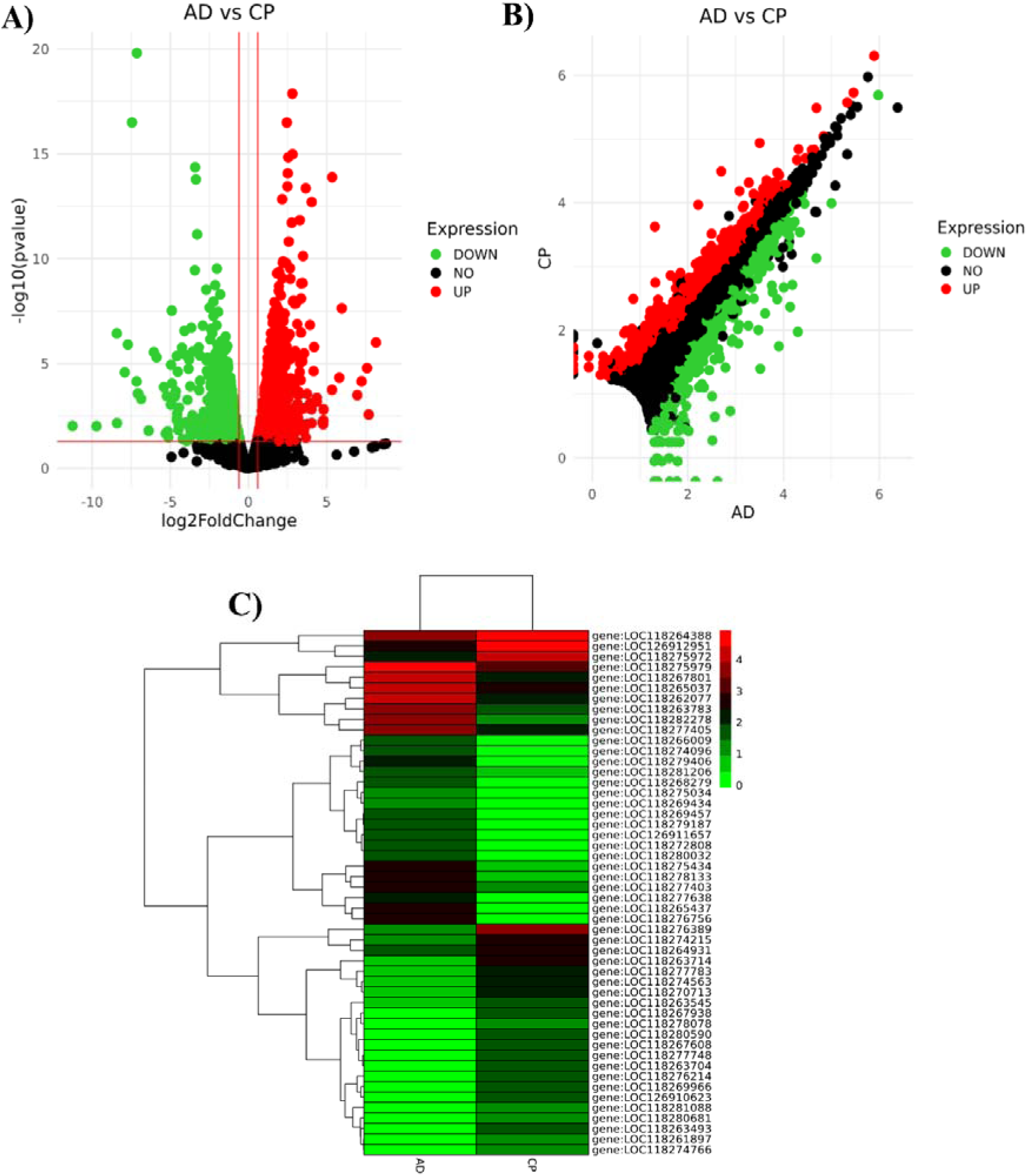
Transcriptome analysis of differentially expressed genes (DEGs) in the salivary glands of *Spodoptera frugiperda* larvae fed on chickpea plants (CP) compared with those fed on artificial diet (AD). **A)** Volcano plot showing the distribution of significantly upregulated and downregulated genes based on log_2_ fold change and statistical significance (−log_10_ *P*-value). Red dots indicate significantly upregulated genes, green dots indicate significantly downregulated genes, and black dots represent genes with no significant differential expression. **B)** Scatter plot showing the overall distribution of gene expression in CP- and AD-fed larvae. Red and green dots represent significantly upregulated and downregulated genes, respectively. **C)** Heat-map depicting the hierarchical clustering and expression patterns of selected DEGs, where red indicates higher expression and green indicates lower expression.

### Gene Ontology & enrichment pathway analysis

Gene Ontology (GO) analysis classified the annotated transcript into three functional categories: biological process, molecular function and cellular component. In the AD-fed versus TP-fed comparison, 14,402 contigs were assigned to 122 GO terms (Supplementary Table S3). These comprised 44 biological-process terms (1,547 genes of 4,188), 47 molecular-function terms (2,355 genes of 5,592), and 31 cellular-component terms (1,237 genes of 4,622) (Fig. 5A).

**Figure 5.**
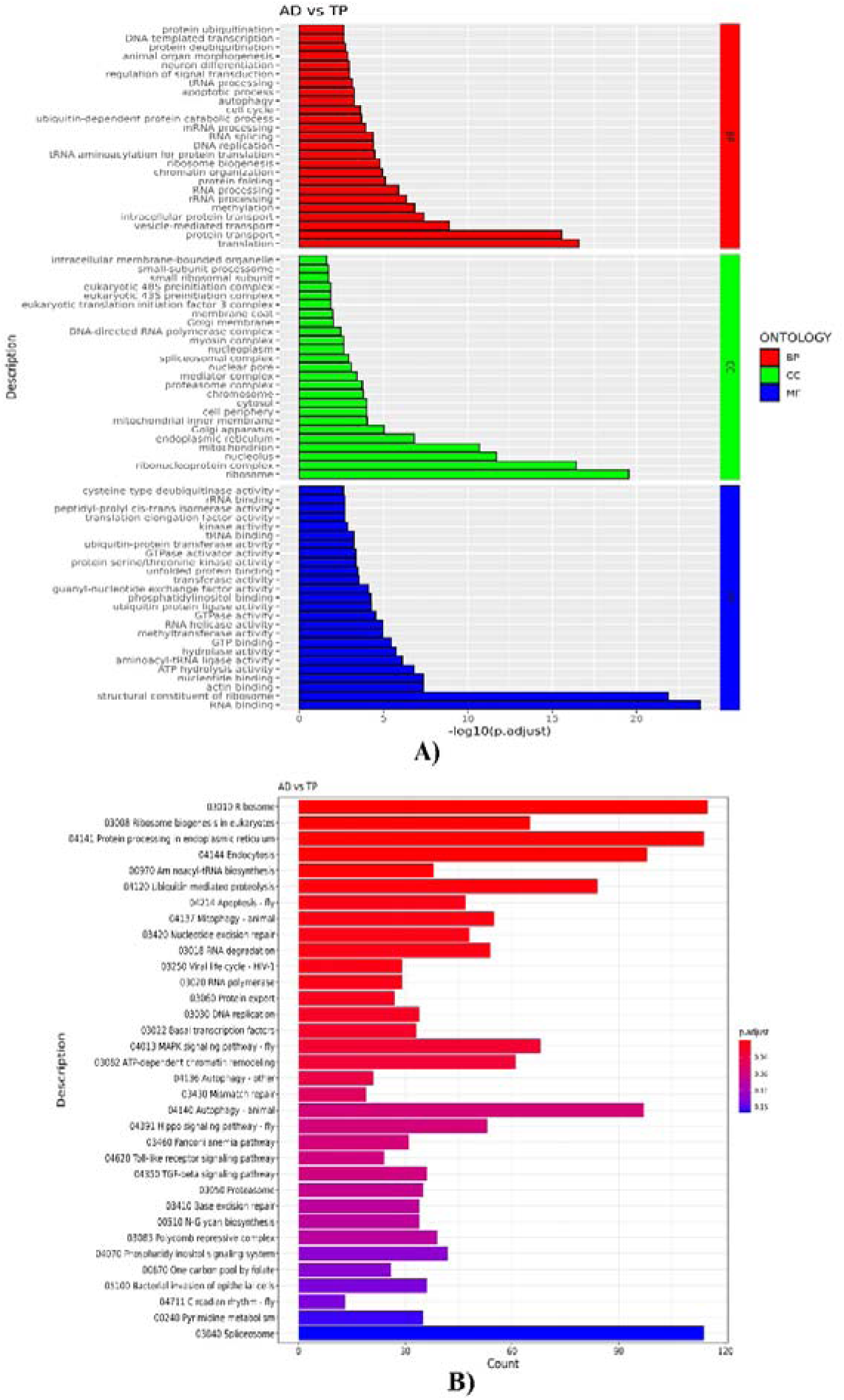
Functional annotation of the transcriptome from the salivary glands of *Spodoptera frugiperda* larvae fed on tomato plant (TP) compared with those fed on artificial diet (AD). **A)** Gene Ontology (GO) classification of differentially expressed genes (DEGs). **B)** KEGG pathway annotation of the identified DEGs.

Similarly, in the AD-fed vs CP-fed comparison, 13,490 contigs were assigned to 127 GO terms (Supplementary Table S4), including 47 biological-process terms (1,542 genes of 3,949), 46 molecular function terms (2,272 genes of 5,208), and 33 cellular component terms (1,255 genes of 4,333) (Fig. 6A).

**Figure 6.**
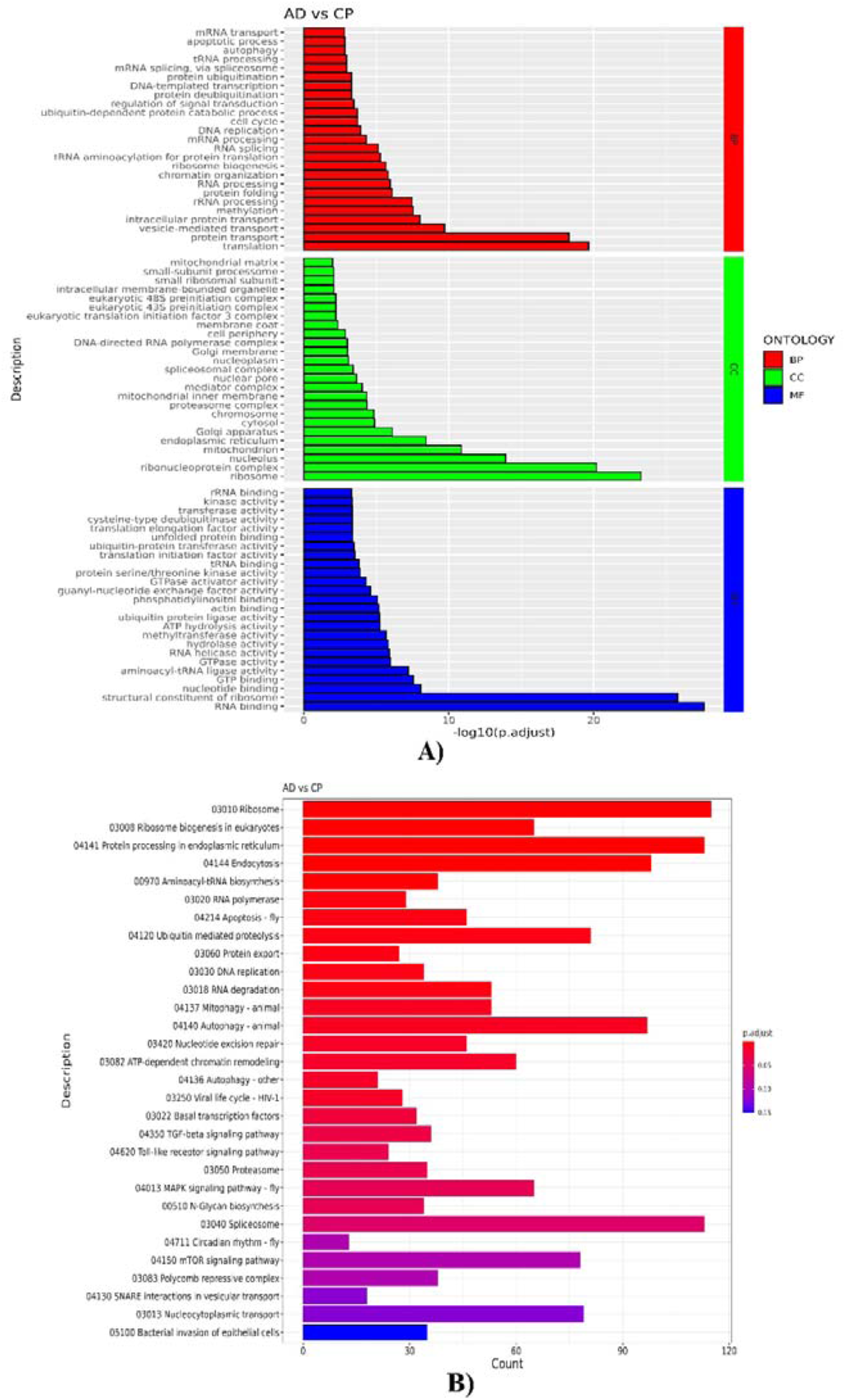
Functional annotation of the transcriptome from the salivary glands of *Spodoptera frugiperda* larvae fed on chickpea plant (CP) compared with those fed on artificial diet (AD). **A)** Gene Ontology (GO) classification of differentially expressed genes (DEGs). **B)** KEGG pathway annotation of the identified DEGs.

Across both comparisons, the predominant biological-process terms included *translation* (GO:0006412), *protein transport* (GO:0015031), and *vesicle-mediated transport* (GO:0016192). The major molecular-function terms included *RNA binding* (GO:0003723) and *hydrolase activity* (GO:0016787), whereas *ribonucleoprotein complex* (GO:1990904), *ribosome* (GO:0005840), and *mitochondrion* (GO:0005739) were among the predominant cellular-component terms.

KASS analysis assigned 3,425 of 14,679 identified genes to KEGG orthologs representing 30 functional categories. The salivary gland transcriptome was mapped to 123 KEGG pathways, of which 34 were significantly enriched (*p* < 0.05) (Supplementary Tables S5 and S6). The most represented enriched pathways were *Ribosome* (115 genes), *Protein processing in the endoplasmic reticulum* (114 genes) and *Spliceosome* (114 genes) (Fig. 5B and 6B).

### Identification of putative secreted proteins

The upregulated DEGs identified in the CP- and TP-fed larvae, relative to AD-fed larvae, were screened to identify diet-associated putatively secreted salivary proteins. A total of 1,017 CP-associated and 353 TP-associated upregulated genes were translated into their corresponding protein sequences and subjected to the *in silico* secretome prediction pipeline.

SignalP predicted signal peptides in 188 CP-associated and 63 TP-associated proteins. Absence of the transmembrane domain was confirmed for 714 CP-associated and 276 TP-associated protein candidates. TargetP analysis further identified 205 CP-associated and 64 TP-associated protein candidates based on their subcellular localization. Finally, WoLF PSORT predicted extracellular localization for 51 CP-associated and 43 TP-associated proteins (Fig. 7A).

**Figure 7.**
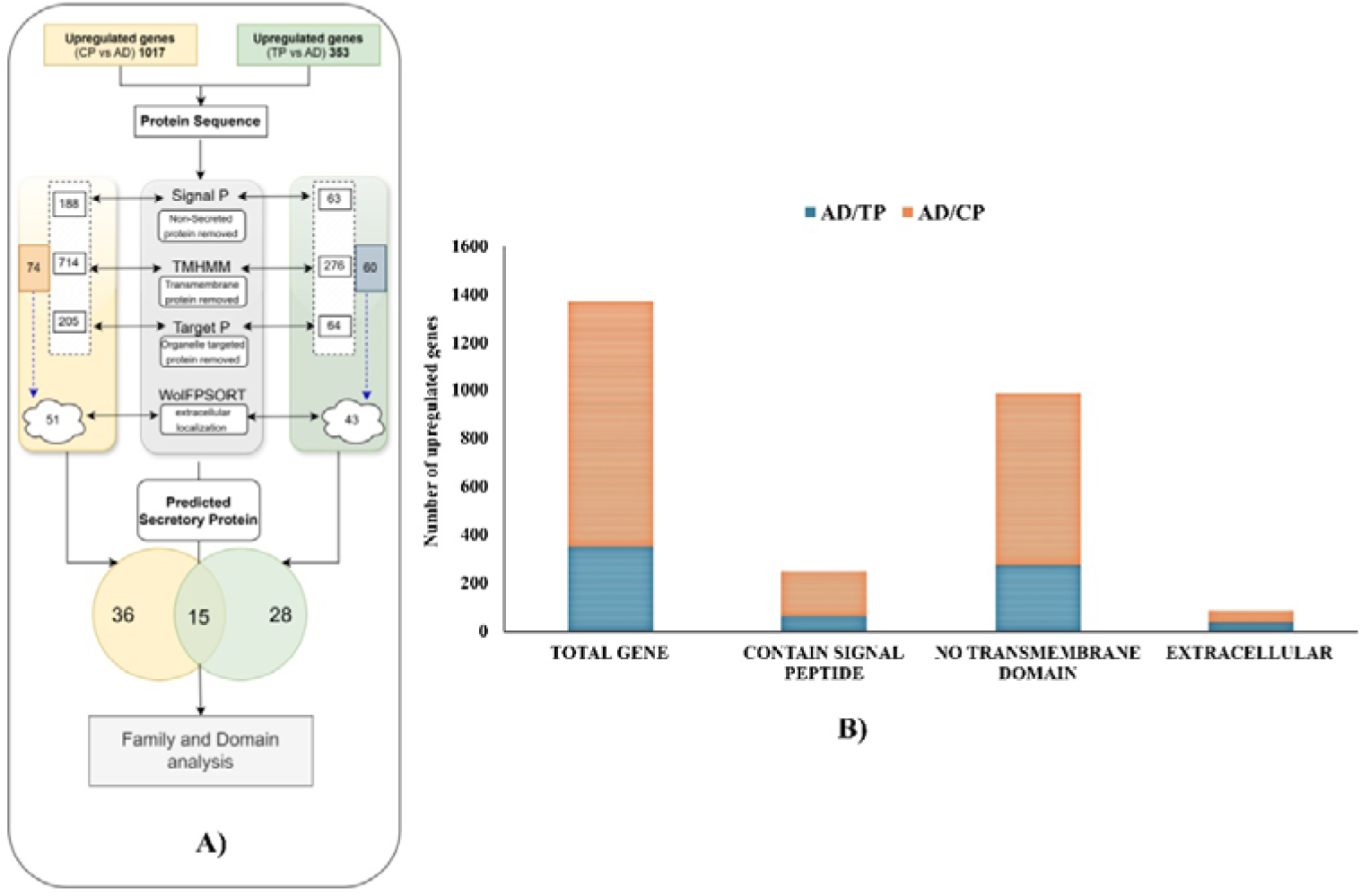
Identification of predicted secretory proteins encoded by upregulated genes in the salivary glands of *Spodoptera frugiperda*. **A)** Upregulated differentially expressed genes (DEGs) from chickpea plant fed (CP; 1017 genes) and tomato plant fed (TP; 353 genes) larvae were screened using Signal P, TMHMM, Target P, and WolFPSORT tools to predict secretory proteins. A total of 51 and 43 putative secretory proteins were identified in the CP and TP datasets, which includes 15 common proteins. The Venn diagram illustrates the overlap between the two datasets, followed by protein family/domain analysis. **B)** Bar graph showing the distribution of secreted proteins containing signal peptide, transmembrane domains, and predicted extracellular localisation signal of genes among the upregulated genes identified in the salivary gland transcriptomes of tomato plant-fed (TP; 354 genes) and chickpea plant-fed (CP; 1,017 genes) *S. frugiperda* larvae.

Thus, 51 proteins from the CP-associated and 43 proteins from the TP-associated salivary transcriptomes fulfilled the criteria for putative secreted proteins (Fig. 7B). Of these, 15 proteins were shared between the two dietary conditions, whereas the remaining candidates were specific to either CP or TP feeding (Supplementary Table S7).

The predicted secretory proteins were further characterized according to protein families, conserved domains and predicted functions (Table 2). A substantial proportion of the predicted secretory proteins in both dietary conditions were uncharacterized. The TP-associated secretome included proteins annotated with structural, developmental and defence-related functions, in addition to uncharacterized proteins, whereas the CP-associated secretome showed a higher representation of structural proteins, enzymes and uncharacterized proteins (Fig. 8).

**Figure 8.**
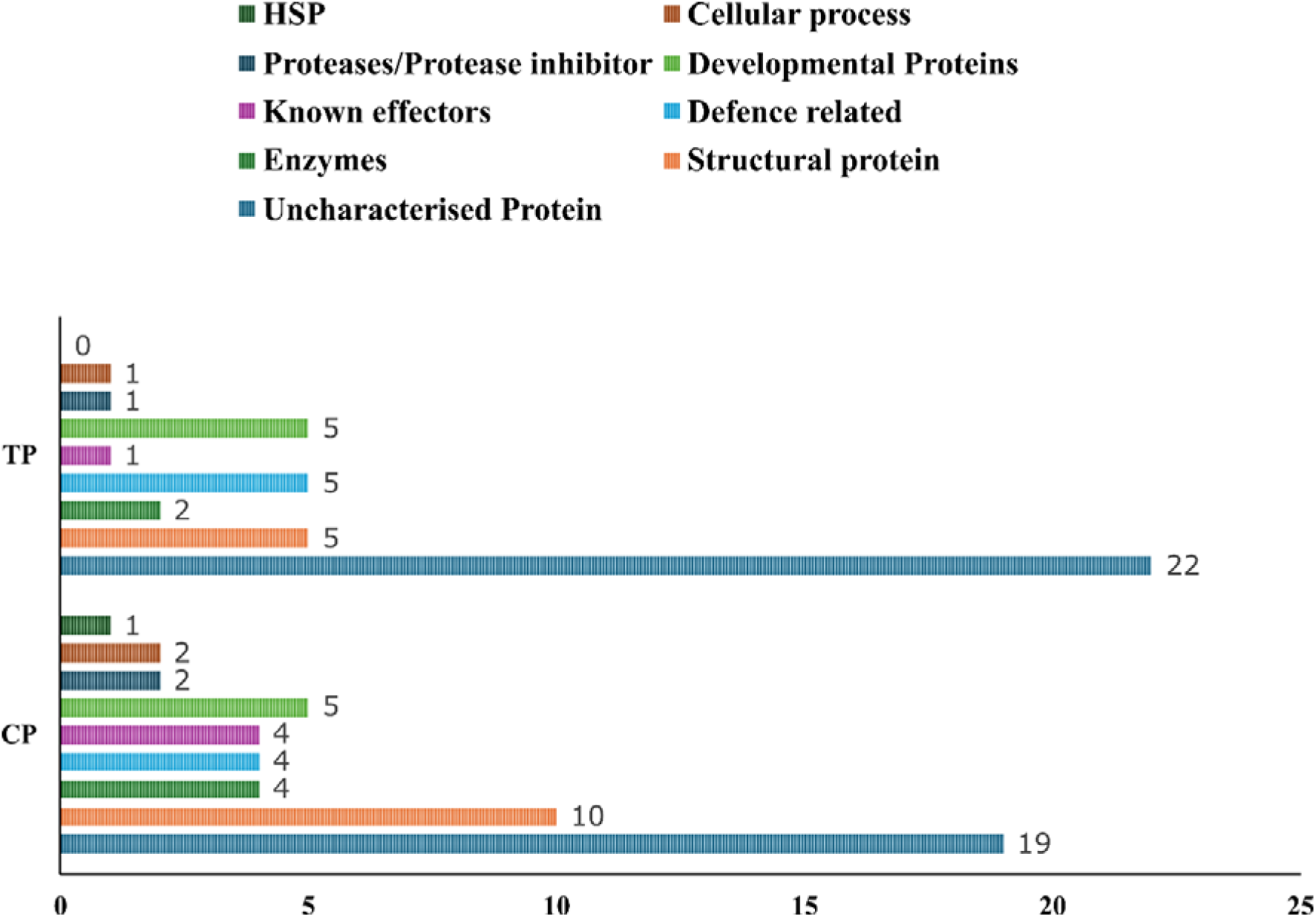
Functional classification of predicted secretory proteins identified in the salivary glands of *S. frugiperda*.

**Table 2:**
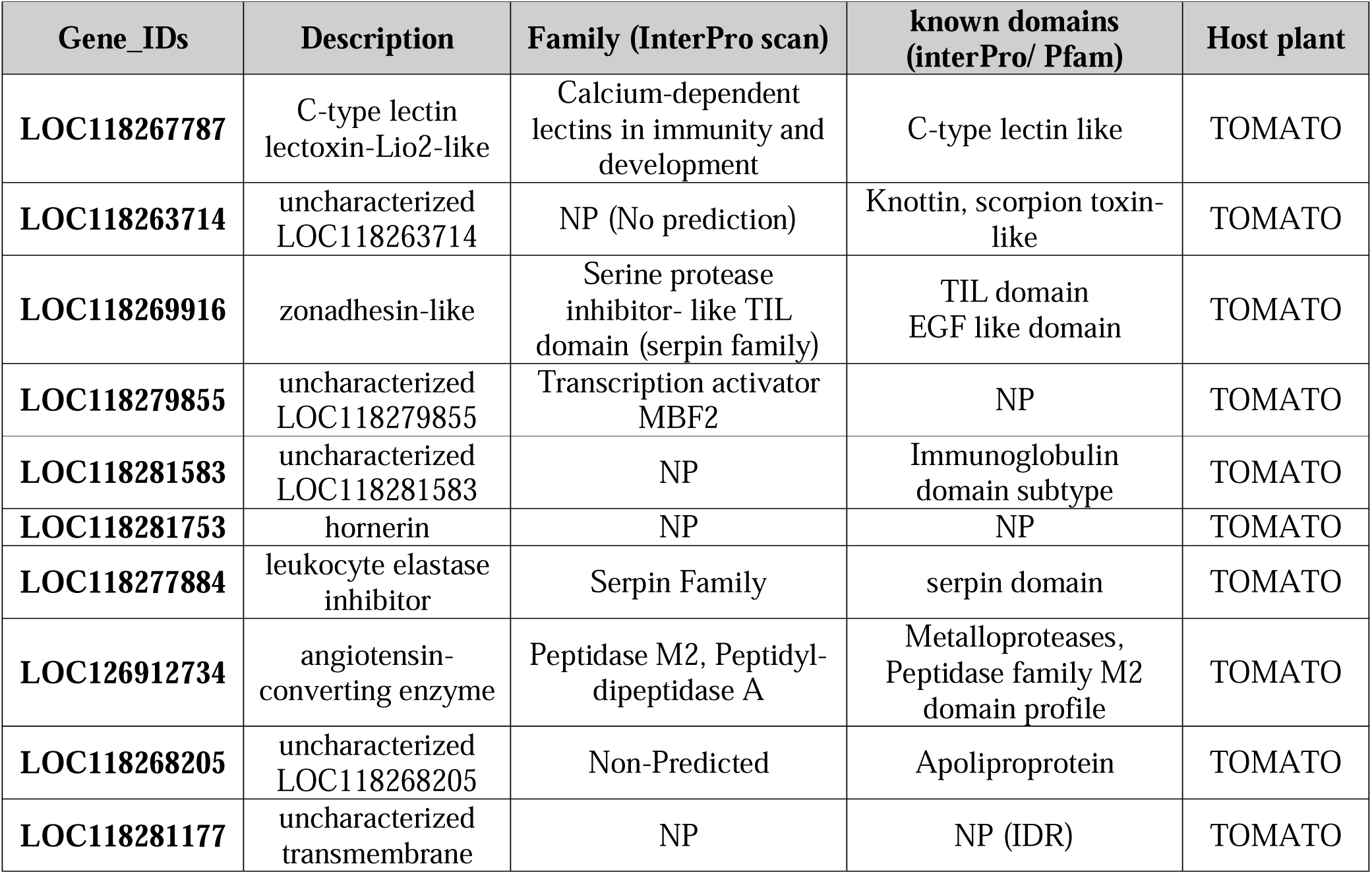
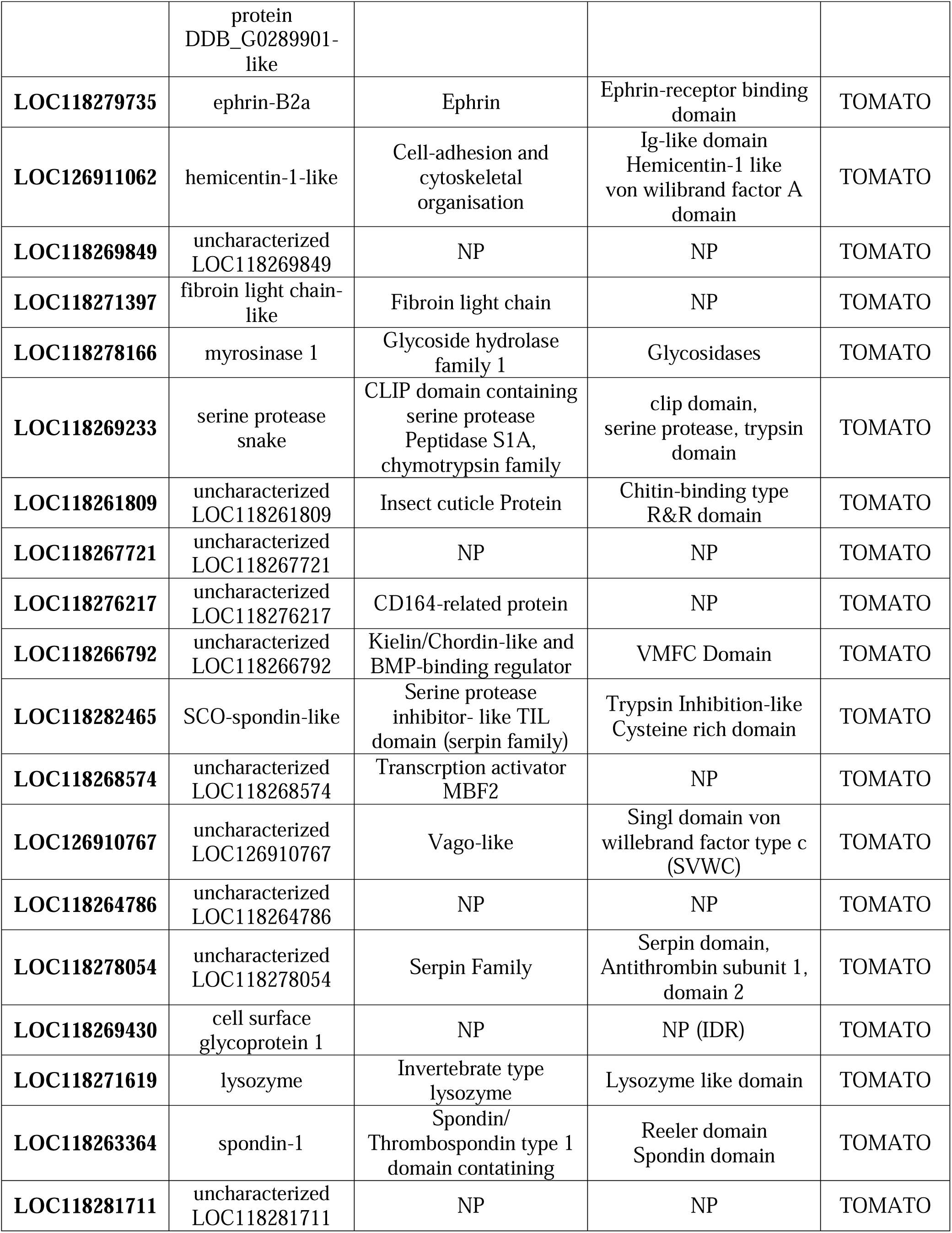
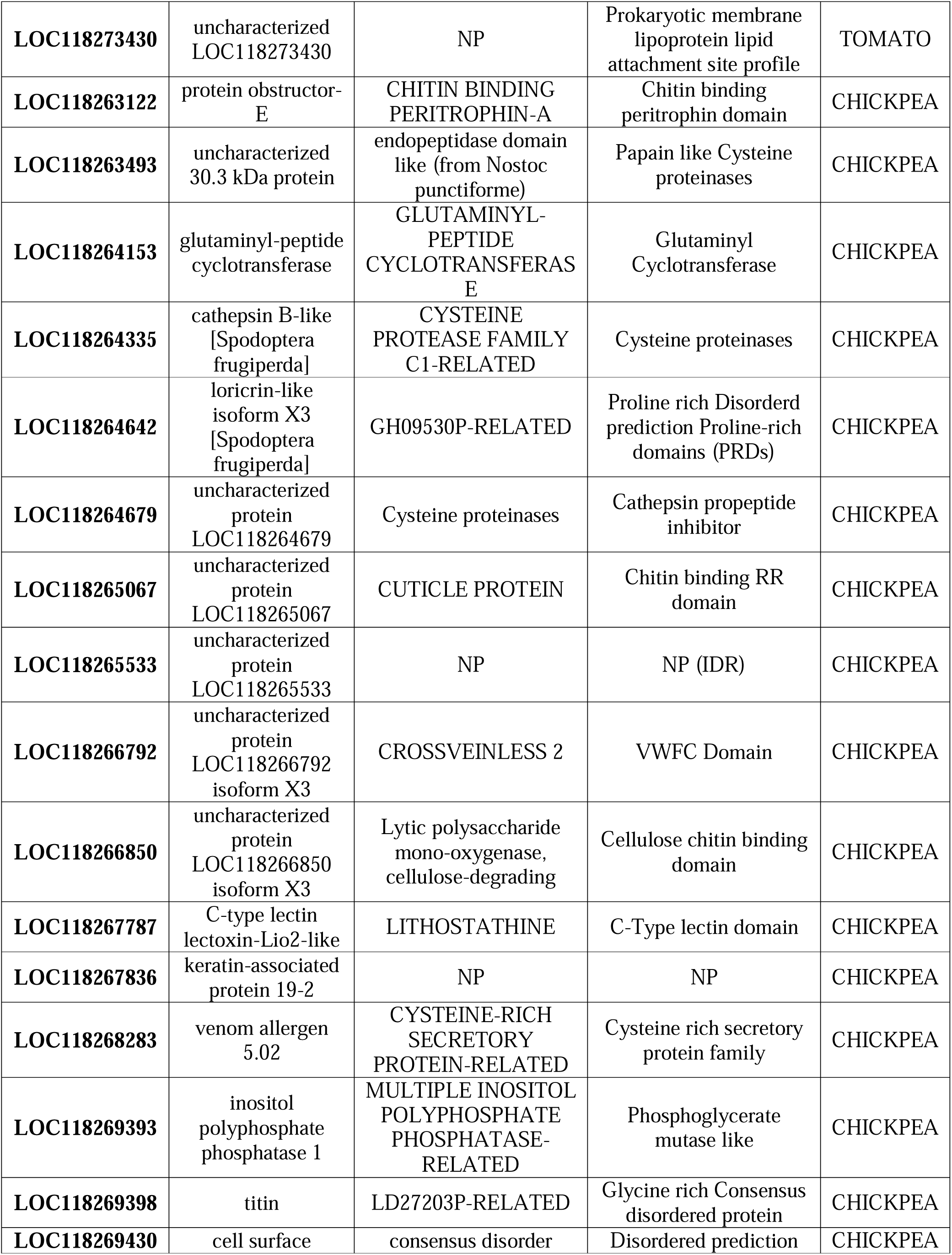
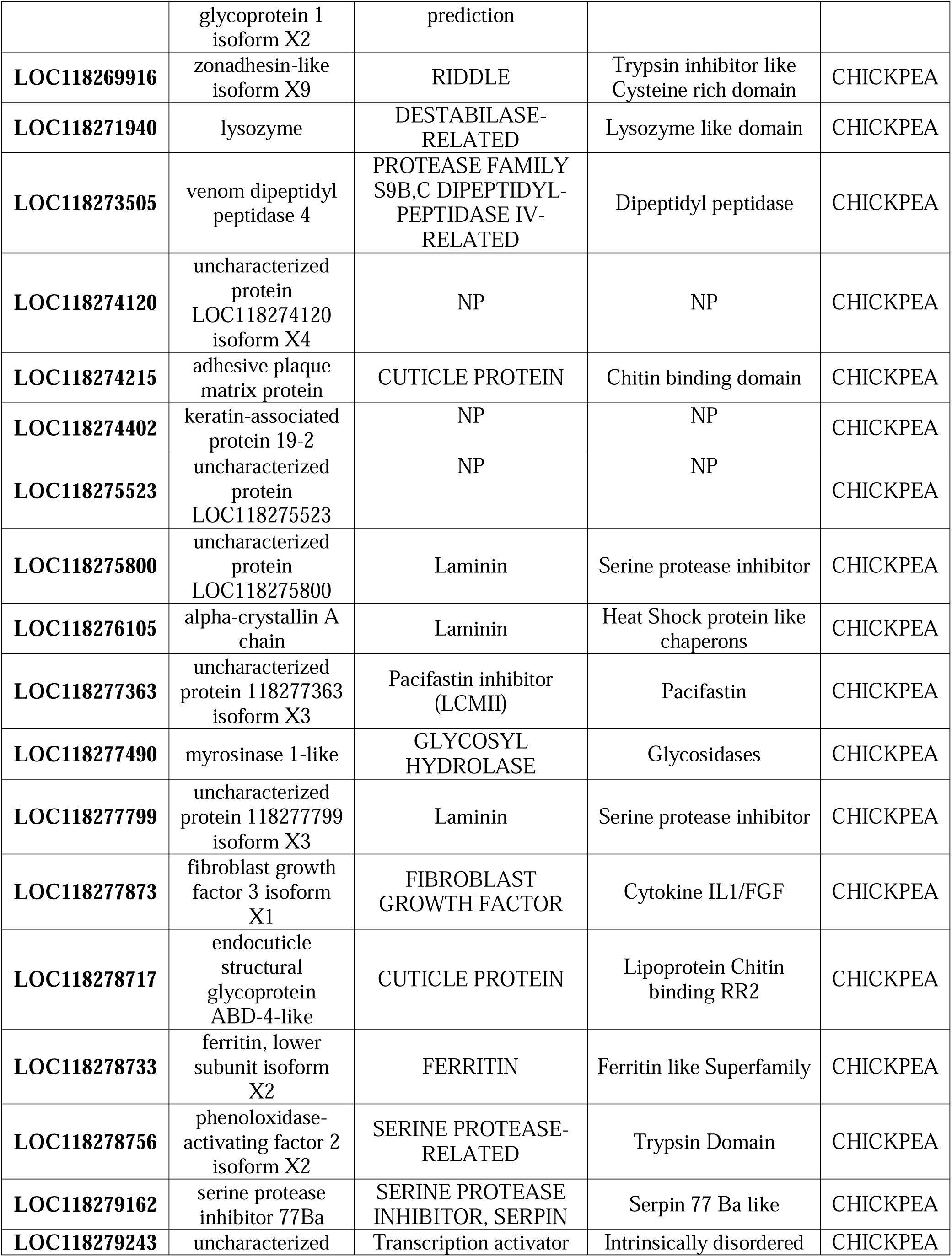
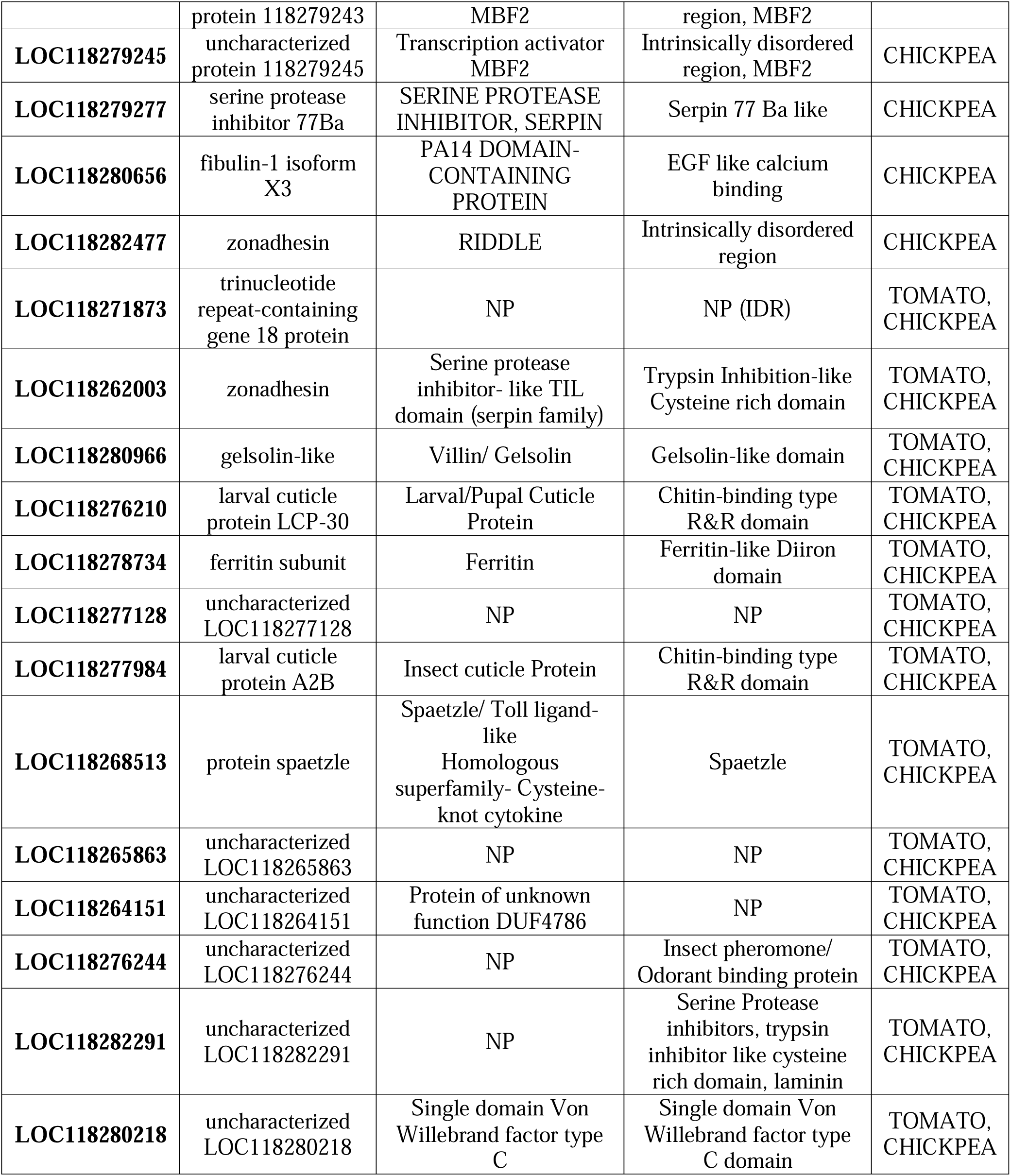
Protein family classification and conserved domains of shortlisted putative secretory proteins identified using InterPro and Pfam.

The physicochemical properties of the shortlisted secretory proteins were assessed using ExPASy ProtParam. The predicted proteins varied substantially in amino acid length, molecular weight, theoretical pI, instability index, aliphatic index and GRAVY value, indicating considerable physicochemical diversity among the candidate salivary proteins (Supplementary Table S8).

### Identification and classification of plant defence-associated proteins

GO annotation-based filtering of the tomato and chickpea reference proteomes identified 55 and 42 biotic stress-responsive proteins, respectively. These proteins were classified into five major defence-related categories: immune regulation, salicylic acid (SA) signalling, jasmonic acid (JA) signalling, reactive oxygen species (ROS) response and insect response (Fig. 9).

**Figure 9.**
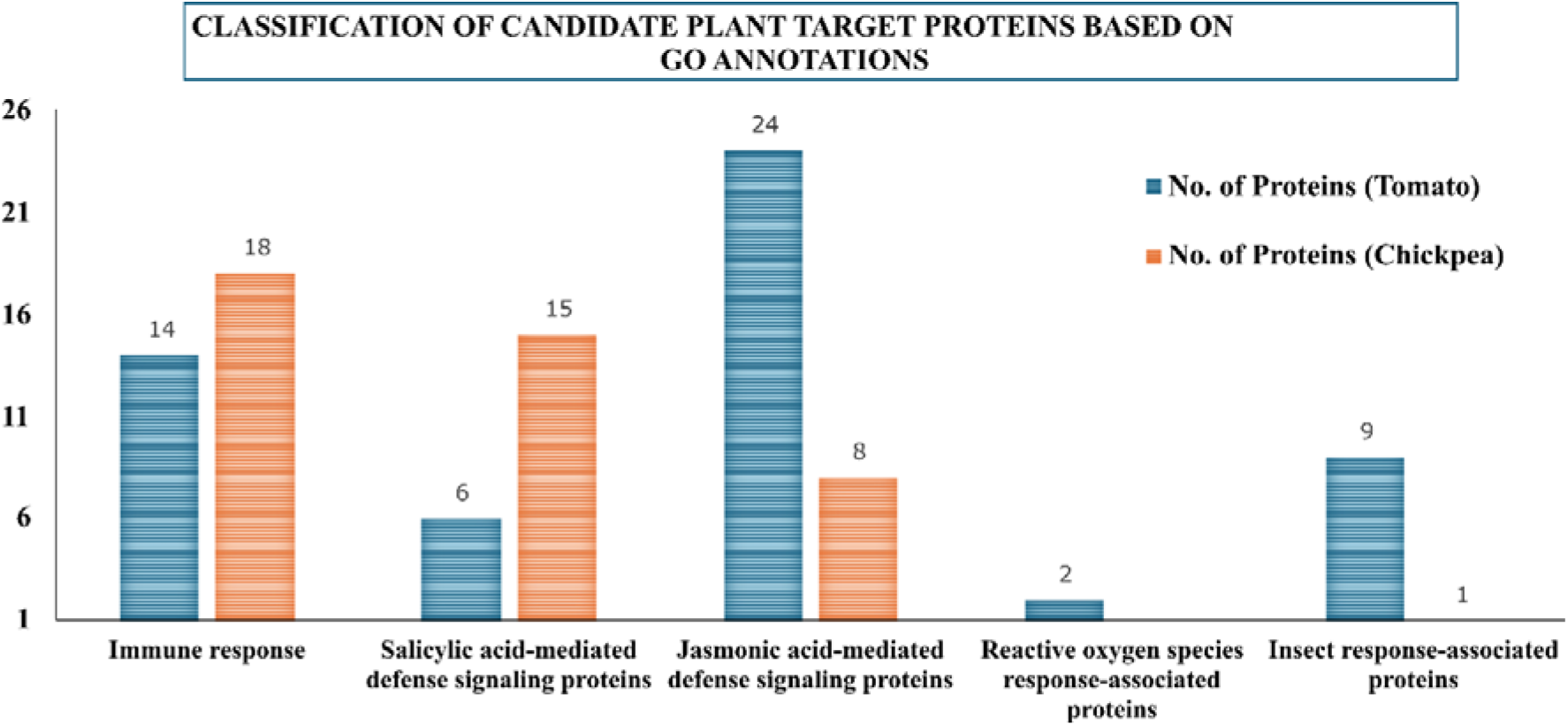
Classification of candidate plant target proteins based on GO annotations in tomato and chickpea plants, identified using the UniProtKB proteome database.

In tomato, the largest group was associated with JA signalling (24 proteins), followed by SA signalling (14), insect response (9), immune regulation (6) and ROS response (2). In chickpea, SA signalling (18 proteins) and immune regulation (15 proteins) represented the largest groups, followed by JA signalling (8) and insect response (1). No chickpea proteins were assigned to the ROS-response category (Fig. 9).

These plant defence-associated proteins were subsequently used as candidate host proteins for protein-protein interaction prediction with the putative secretory proteins of *S. frugiperda*.

### Prediction of putative interactions between salivary proteins and plant defence-associated proteins

The 51 CP-associated and 43 TP-associated putative secretory proteins were screened against 42 chickpea and 55 tomato defence-associated proteins, respectively, using AlphaPulldown. This resulted in 2,142 predicted protein-protein interaction pairs for chickpea and 2,365 interactions for tomato (Supplementary Tables S9 and S10).

The predicted interactions were ranked according to interface predicted TM-score (iPTM). The top 10 insect secretory protein-plant protein pairs from each dataset were selected using an iPTM threshold of >0.7. PAE and pLDDT values were additionally considered to assess the confidence of the predicted interactions and structural models.

The highest-confidence interaction pairs were re-examined after removal of the predicted N-terminal signal peptide from the secretory proteins, reflecting the expected processing of proteins during secretion. Heat maps based on iPTM scores were generated for the top interaction candidates from the TP and CP datasets (Figs. 10 and 11).

**Figure 10.**
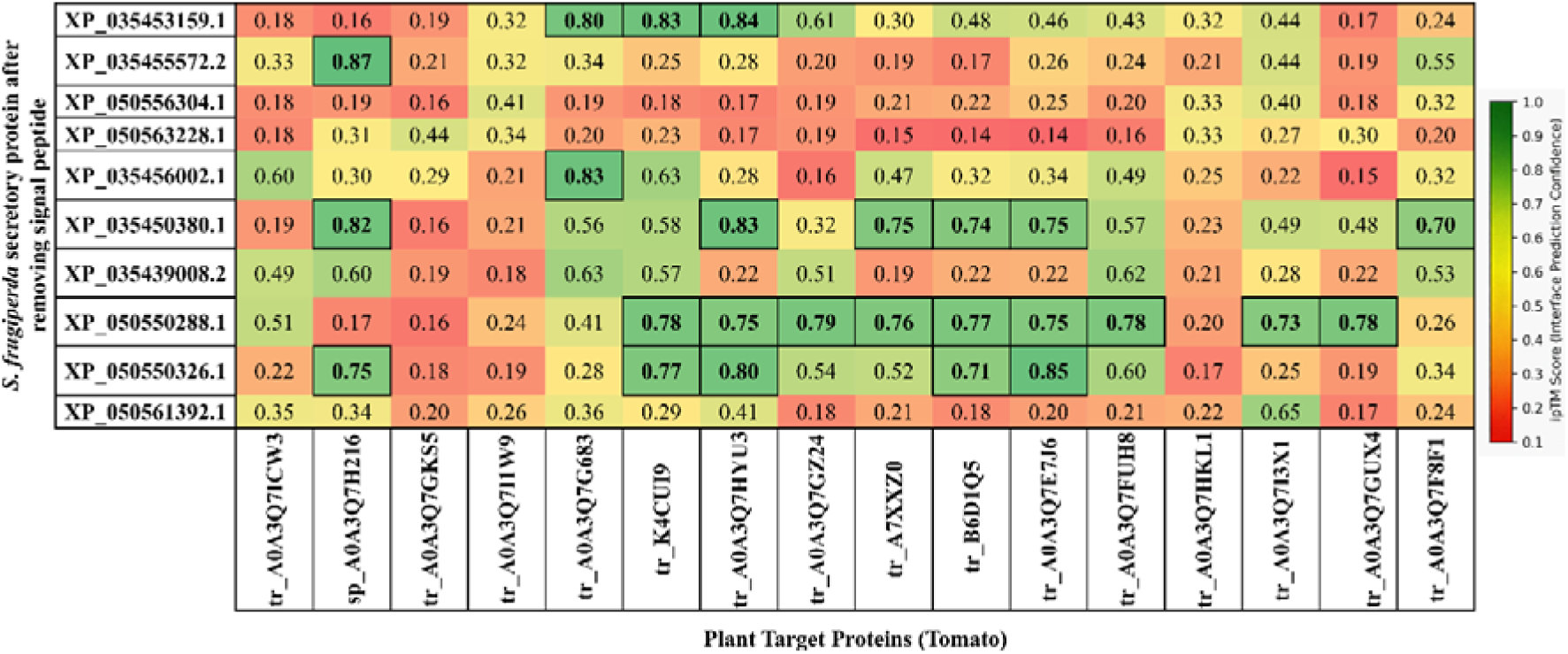
Heat-map illustrating top predicted interactions between *S. frugiperda* secretory proteins (without signal peptide) and tomato plant target proteins using AlphaPulldown v2.1.1.

**Figure 11.**
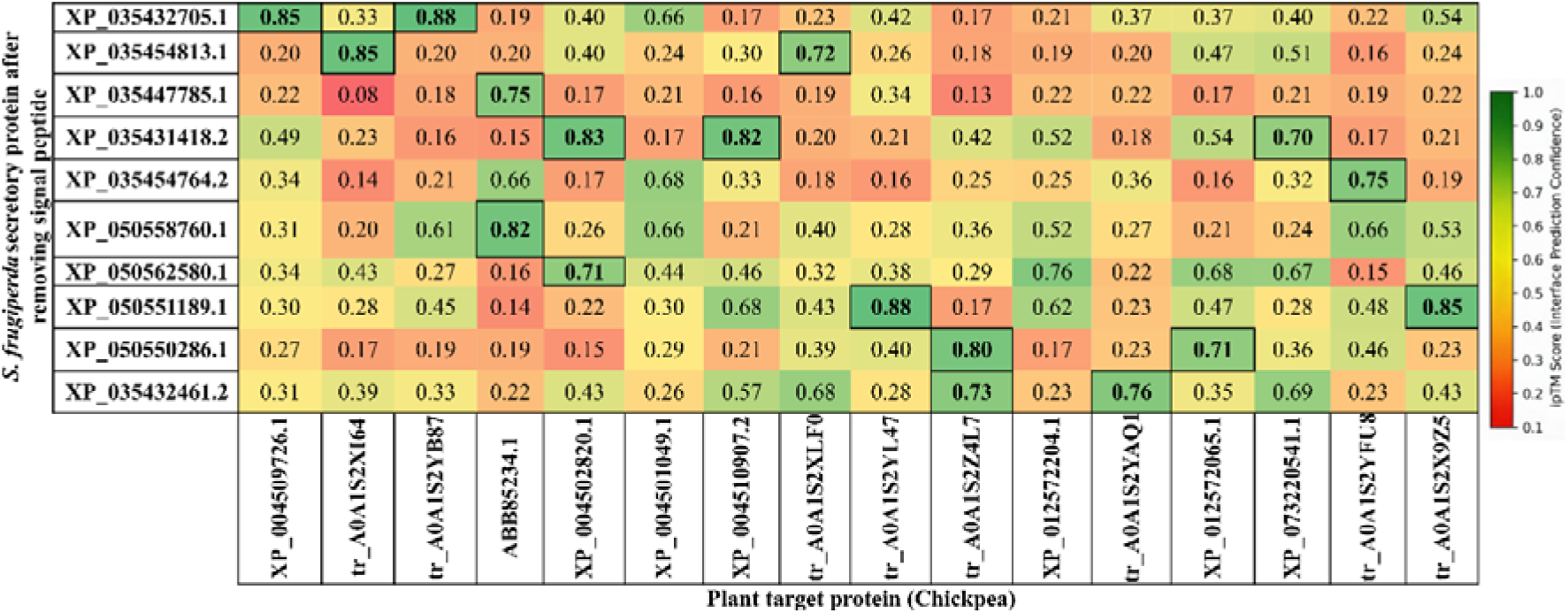
Heat-map illustrating top predicted interactions between *S. frugiperda* secretory proteins (without signal peptide) and chickpea plant target proteins using AlphaPulldown v2.1.1.

Comparative analysis showed that removal of the signal peptide increased the predicted interaction confidence for several protein pairs. Representative high-confidence interactions were further evaluated using PAE plots and predicted complex structures for both full-length and signal-peptide-trimmed forms of the *S. frugiperda* proteins (Figs. 12 and 13). Removal of the N-terminal signal peptide did not substantially alter the predicted interaction interfaces but resulted in increased iPTM scores for the evaluated complexes.

**Figure 12.**
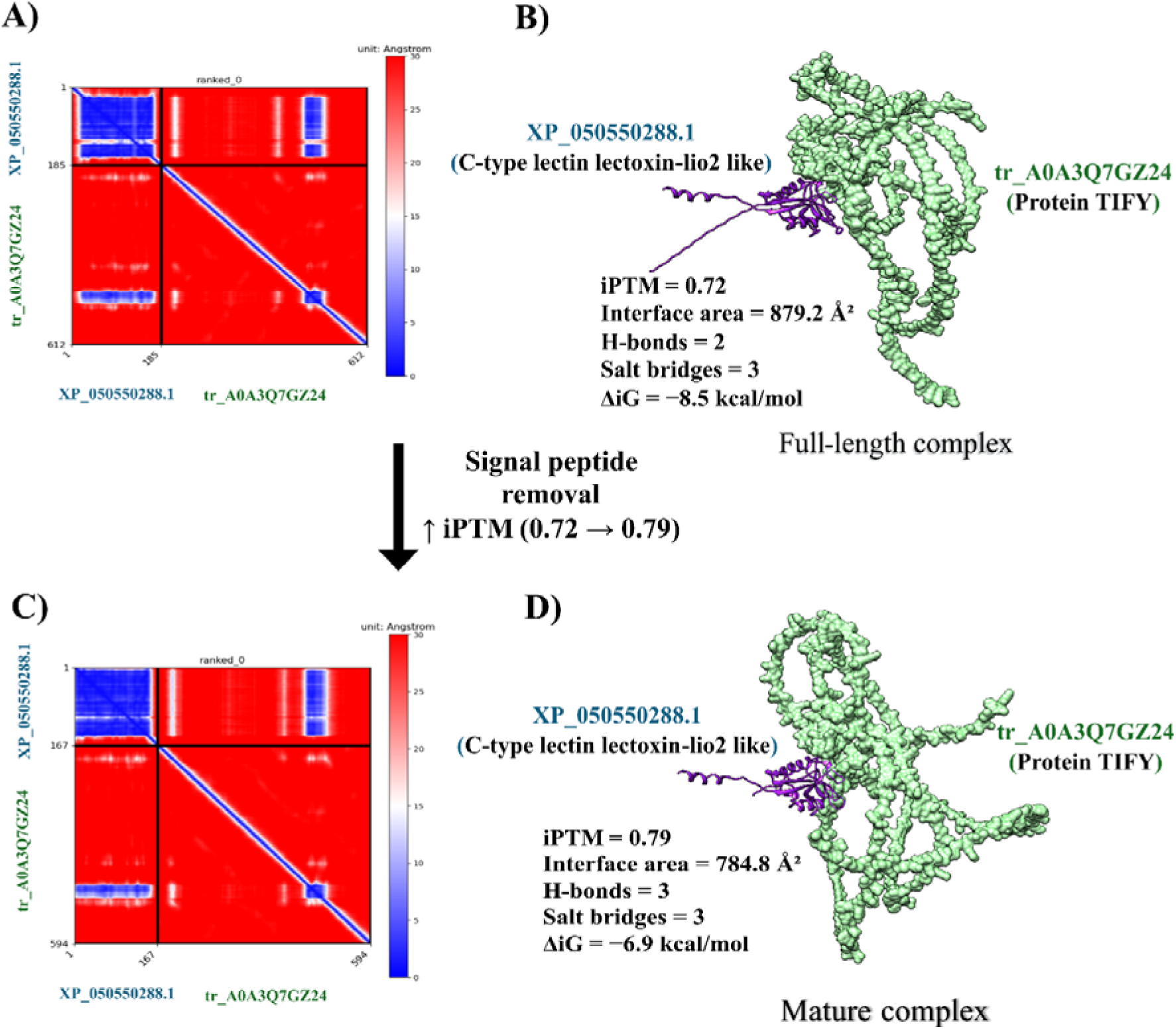
Comparison of AlphaPulldown predicted interactions between full-length and mature forms (without signal peptide) of *S. frugiperda* secretory protein, XP_050550288.1 (C-type lectin lectoxin-lio2 like) and tomato plant target protein, tr_A0A3Q7GZ24 (Protein TIFY). **A) and C)** PAE plots of the full-length and mature protein complexes, respectively. **B) and D)** Corresponding predicted protein structures.

**Figure 13.**
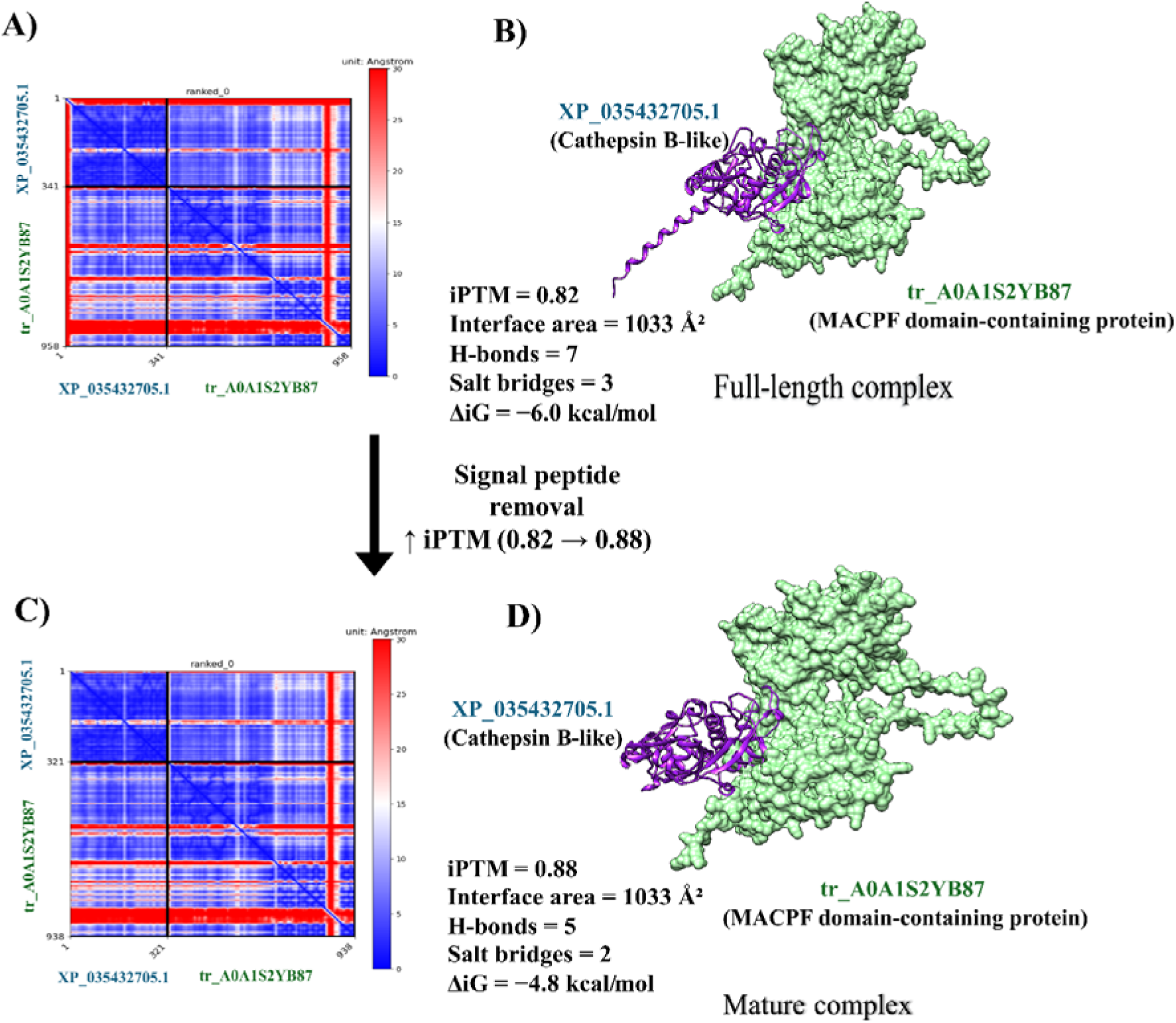
Comparison of AlphaPulldown prediction interactions between full-length and mature forms (without signal peptide) of *S. frugiperda* secretory protein, XP_035432705.1 (Cathepsin B-like) and chickpea plant target protein, tr_A0A1S2YB87 (MACPF domain- containing protein). **A) and C)** PAE plots of the full-length and mature protein complexes, respectively. **B) and D)** Corresponding predicted protein structures.

## Discussion

*Spodoptera frugiperda* is a polyphagous herbivore whose growth, development, and overall fitness vary considerably among host plants. Previous studies have shown greater larval growth, survival, and reproductive fitness on preferred hosts such as maize than on alternative hosts, including tomato and chickpea (Putra, Dyati, & Afrindini, 2024; Shoman, Ghanim, Harraz, & Aziz, 2025). Here, we examined host-associated changes in the salivary gland transcriptome of *S. frugiperda* larvae feeding on tomato and chickpea compared with an artificial diet, with particular emphasis on putatively secreted proteins and their predicted interactions with plant defence-associated proteins.

Both, salivary gland transcriptomics and salivary secretion proteomics have been used to identify candidate insect effectors involved in host adaptation (Carolan et al., 2011). We focused on salivary gland transcriptomics because transcript-based approaches can capture candidate secreted proteins that may be underrepresented or undetectable in saliva proteomes. For example, Bt56 and LsSP1 were detected through salivary gland transcriptomics but not in secreted saliva proteomic analyses (Huang et al., 2021; X. J. Wang, Li, Ye, & Huang, 2024).

A notable finding was the larger number of DEGs in chickpea-fed larvae than in tomato-fed larvae, indicating a stronger transcriptional response to chickpea feeding (Fig. 1; Supplementary Tables S1 and S2). The greater number of differentially expressed transcripts was accompanied by a higher number of predicted secretory proteins in chickpea-fed larvae (Fig. 7), suggesting that feeding on chickpea is associated with broader transcriptional and secretory responses. This broader response may reflect the need to cope with host-specific chemical or anti-herbivore defences, although the present data do not establish the physiological basis of this response.

A large proportion of the predicted secretory proteins were unannotated; however, several contained conserved domains, including immunoglobulin-like, von Willebrand factor type C (VWFC), MBF2-related, and knottin-like domains. Immunoglobulin-like proteins have been associated with pathogen recognition and innate immune defence in insects (Li et al., 2025; Mandrioli, Monti, & Tedeschi, 2015), while VWFC-containing proteins have been implicated in insect immune and stress responses (Y. Chen et al., 2026; Labropoulou, Wang, Magkrioti, Smagghe, & Swevers, 2024). The occurrence of these domains in host-responsive secreted proteins highlights their potential functional relevance, but their roles in plant–insect interactions remain to be established.

Of particular interest were MBF2-related proteins, characteristic of insect REPAT proteins. REPAT proteins are associated with stress and immune responses in lepidopteran insects (Navarro-Cerrillo, Hernández-Martínez, Vogel, Ferré, & Herrero, 2013), and MBF2- containing REPAT proteins have recently been implicated in suppression of plant defence in *Phthorimaea absoluta* and *Spodoptera* species (Wang et al. 2024a) (García-Marín et al. 2025). Moreover, REPAT38 from *Spodoptera exigua* shares high sequence identity with HARP1 from *Helicoverpa armigera*, which interacts with Arabidopsis JAZ proteins and modulates jasmonate signalling (C. Y. Chen et al., 2019). These observations support the REPAT proteins identified here as candidates for further investigation as salivary effectors, although their effector activity in *S. frugiperda* remains unverified.

Other predicted secreted proteins included Spaetzle, lysozymes, C-type lectin, cathepsin B- like protein, serine protease inhibitors, venom allergen 5, and venom dipeptidyl peptidase 4 (Supplementary Table S1). Their known roles in immunity, proteolysis, and host interaction provide plausible functional links to salivary-mediated host adaptation. Chickpea-fed larvae showed a higher abundance of proteases and protease inhibitors, including cathepsin B-like protein and serine protease inhibitors (Table 2), which may reflect responses to plant anti- digestive defences. Salivary cathepsin B has been reported in *Myzus persicae* and *Nilaparvata lugens*, and proteases can counter plant protease inhibitors by degrading these anti-digestive proteins (Chesnais et al., 2022; Foissac, Edwards, Du, Gatehouse, & Gatehouse, 2002) (Zhu-Salzman & Zeng, 2015). The presence of venom-associated proteins further expands the repertoire of candidate host-interacting proteins, although their specific functions in *S. frugiperda* feeding remain to be determined.

Predicted protein–protein interactions provided an additional layer for prioritizing candidate effector-host target relationships. AlphaPulldown identified several secreted protein-plant defence protein pairs with high predicted interaction scores (Figs. 10 and 11; Supplementary Tables S9 and S10). Removal of N-terminal signal peptides increased the predicted interaction confidence for several pairs, consistent with the possibility that signal peptides can influence structural modelling of mature secreted proteins. Similar approaches have been used in structural studies of secreted effectors (Seong and Krasileva 2023) (Sanaboyana and Elcock 2024). However, signal peptide removal did not improve all interactions, indicating that this step should be considered as a refinement rather than evidence of biological interaction.

One of the highest-confidence predicted interactions involved a C-type lectin lectoxin-lio2- like protein and a tomato TIFY family protein (Fig. 12). TIFY proteins, particularly JAZ proteins, are important regulators of jasmonic acid signalling and plant defence against herbivores (X. Zhao, He, Liu, Wang, & Zhao, 2024). The involvement of *SlJAZ* genes in tomato responses to *Tuta absoluta* infestation further supports the relevance of this pathway to insect–plant interactions (Haq et al., 2025). C-type lectins are traditionally associated with innate immune recognition in insects but have also been implicated in host manipulation in parasitic nematodes (J. Zhao et al., 2021; Zhuo et al., 2019) (Chi et al., 2024). The predicted interaction therefore raises the possibility that the *S. frugiperda* C-type lectin may influence JA-associated defence signalling through interaction with a tomato TIFY protein. This interpretation remains hypothetical and requires experimental validation.

Another high-confidence interaction involved a *S. frugiperda* cathepsin B-like protein and a chickpea MACPF domain-containing protein (Fig. 13). Plant MACPF proteins have been associated with immune responses, stress signalling, and programmed cell death (Ma et al., 2023; Xue Zhang et al., 2022), although their specific role in defence against herbivorous insects remains unclear. Cathepsin B is primarily recognized as a cysteine protease but has also been implicated in modulation of plant immunity. For example, aphid-derived cathepsin B proteins can target defence regulators including EDS1, PAD4, and ADR1 (Liu et al., 2025). The predicted cathepsin B–MACPF interaction therefore identifies a potentially important host–salivary protein pair for future investigation. These findings demonstrate that insect-derived cathepsin B proteins function not only in digestion but also as effectors that directly manipulate plant defence pathways. Whether the *S. frugiperda* cathepsin B-like protein similarly modulates chickpea defence through interaction with MACPF proteins remains to be determined experimentally.

In conclusion, this study reveals pronounced host-associated changes in the salivary gland transcriptome of *S. frugiperda* and identifies a set of host-responsive putatively secreted proteins, including REPAT, C-type lectin, cathepsin B-like, and venom-related proteins. Integration of secretome prediction with protein-structure-based interaction modelling further identified candidate plant defence-associated targets, including TIFY and MACPF proteins. These candidates provide testable hypotheses for understanding how salivary proteins may contribute to host adaptation and manipulation of plant defence. Functional studies will be required to determine whether these predicted proteins and interactions contribute directly to host defence suppression or feeding adaptation.

## CRediT authorship contribution statement

**Sakshi Pandey:** Writing – original draft, Investigation, Methodology, Formal analysis, validation. **Sundaram Shilpi:** Writing – original draft, Investigation, Methodology, Formal analysis, validation. **Vineeta Roshanlal**: Investigation, methodology**. Hanuman Meena:** Investigation, methodology. **Chandra Pal Singh:** Investigation, review & editing, Supervision. **Jayendra Nath Shukla:** Investigation, Validation, review & editing, Supervision, Resources, Project administration, Conceptualization.

## Declaration of competing interest

The authors declare no conflict of interests.

## Ethical Approval

Not applicable

## Funding sources

This work was supported by the grant (CRG/2023/006907) from the Department of Science and Technology, Ministry of Science and Technology, India.

## Acknowledgements

1. J. N. Shukla gratefully acknowledge the Ramalingaswami Re-entry Fellowships (BT/RLF/Re- entry/10/2015) awarded by the Department of Biotechnology, New Delhi. JNS also acknowledges the financial supports through the DST SERB research grants (CRG/2023/006907 and EMR/2017/001378/AS), New Delhi, and the DBT Builder project (BT/INF/22/SP44383/2021) for providing workstation facilities that supported this work.

## Data availability

The raw RNA-seq data generated during this study have been deposited in the NCBI Sequence Read Archive (SRA) under BioProject accession **PRJNA1496506** (SRA accessions: **SRR39776117–SRR39776122**).

## Notes

### Competing Interest Statement

The authors have declared no competing interest.

